# Clustering laminar cytoarchitecture: in vivo parcellation based on cortical granularity

**DOI:** 10.1101/2022.10.29.514347

**Authors:** Ittai Shamir, Yaniv Assaf, Ron Shamir

## Abstract

The laminar microstructure of the cerebral cortex is considered a unique anatomical mark of the development, function, connectivity, and even various pathologies of the brain. In recent years, multiple neuroimaging studies have utilized magnetic resonance imaging (MRI) relaxometry to visualize and explore this intricate microstructure. By successfully delineating the cortical laminar components, the applicability of T1 relaxometry has been expanded beyond solely a direct measure of myeline content. However, validating its applicability as a measure of cortical cytoarchitecture demands analyzing the complex resulting laminar datasets. In this study, we adapt and implement an algorithm for clustering cell omics profiles to cluster these complex microstructural cortical datasets. The resulting clusters correspond with an established atlas of cytoarchitectonic features, providing robust validation of T1 imaging as a tool for exploring cortical laminar composition. Lastly, we demonstrate the applicability of this framework in the exploration of the cytoarchitectonic features behind various unique skillsets.

## 1. Introduction

### 1.1 Progress in MRI imaging of T1 layers

The intricate laminar structure of the cerebral cortex was first discovered in the beginning of the twentieth century using ex-vivo histological methods, sparking a century of studies into its assumed roles in the development, function, connectivity, and even pathologies of the brain. With the advent of MRI neuroimaging, the cerebral cortex was successfully segmented, delineating its cortical surfaces bordering with underlying white matter and the surrounding cerebrospinal fluid (Fischl, 2012). However, until about a decade ago the laminar substructure of the cerebral cortex was assumed to be beyond the imaging capabilities of MRI. Since then, multiple neuroimaging studies have proposed a variety of MRI imaging modalities and approaches for exploring the laminar composition of the cortex, with T1 relaxometry proving to be the most suitable and accurate approach so far.

The applicability of T1 relaxometry in exploring the cortical laminar composition was established thanks to a series of studies. In 2012, a 2012 study characterized the cortical layers in the brains of both humans and rats and compared the resulting T1 clusters to histological findings from the rat brain (Barazany and Assaf, 2012). In 2018, a larger scale study used the same inversion recovery (IR) MRI protocol to explore the laminar composition of both humans and rats, concluding that low resolution T1 mapping is the most appropriate approach for delineating the layers (Lifshits et al., 2018). In 2019, a complete automated framework was formed for analyzing the cortical laminar composition using low resolution multi-T1 mapping and a simple surface-based volumetric sampling system (Shamir et al., 2019). What followed were several studies using multi-T1 imaging to explore the role of the cortical laminar composition in different pathologies, such as epilepsy (Lotan et al., 2021), as well as in healthy aging (Tomer et al., 2022). Other studies modeled and explored patterns of cortical connectivity on the laminar level (Shamir and Assaf, 2021a, 2021b; Shamir et al., 2022).

The abovementioned studies provide further validation for cortical laminar composition analysis using low resolution, multi-T1 imaging. However, this methodology is still limited by the fact that T1 is not considered a direct measure of cytoarchitecture. Despite the established correspondence between T1 layers and the actual cortical layers, T1 is still considered more of a direct measure of myeloarchitecture (myeline content) than of cytoarchitecture (cellular composition).

### 1.2 Challenges in clustering multilayered surface-based data

Use of the framework for cortical laminar composition analysis (Shamir et al., 2019; Shamir and Assaf, 2021a, 2021b; Shamir et al., 2022) results in a multilayered, surface-based dataset representing the regionally varying laminar composition across the cortical surfaces of both hemispheres. This dataset is both multidimensional and geometrically complex: six cortical laminar components, representing the regionally varying microstructure of the cortex, are measured across vertices of a Delaunay triangulation, delineating the intricate geometry of the cortex. More specifically, for each hemisphere the triangulation consists of ~150,000 vertices, connected by ~300,000 faces, with laminar composition values available for each vertex, including six laminar components corresponding to the widths of T1 layers 1-6 (see figure 1, part 1).

**Figure 1:**
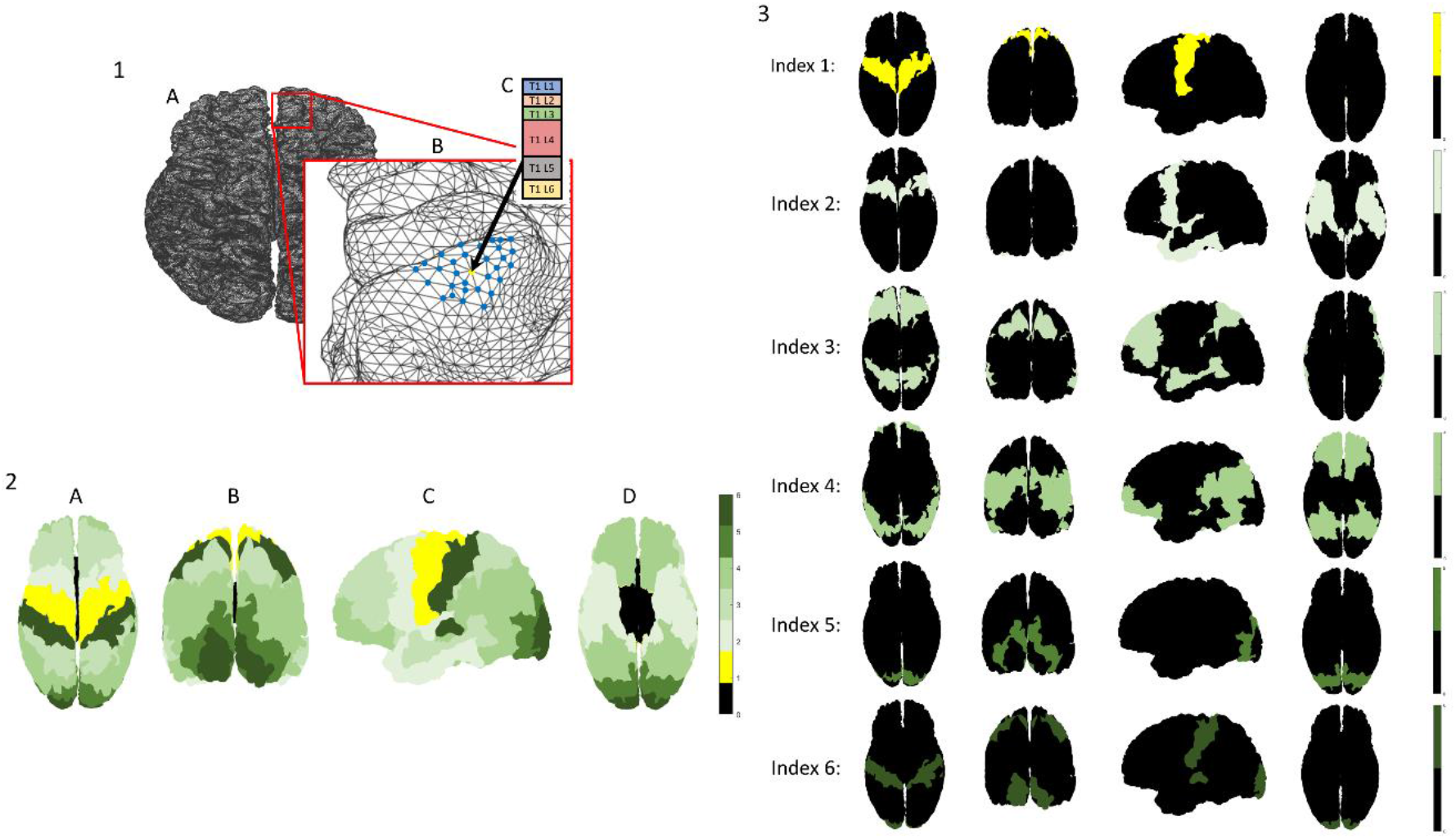
Multilayered surface-based dataset and a cytoarchitectonic granularity atlas: 1- Multilayered cytoarchitectonic surface-based dataset: the cerebral cortex is represented by a Delaunay triangulation, delineating the mid cortical surfaces of each hemisphere (seen from a top view in A). A single vertex is seen (yellow) on a sulcus in the frontal lobe of the right hemisphere, surrounded by its thirty closest neighboring vertices (blue) (B). The cytoarchitectonic laminar composition at the location of the chosen vertex is shown, including widths of six laminar components: T1 layers 1-6 (colored individually in C) 2-3- A cytoarchitectonic atlas of granularity indices: cytoarchitectonic labelling of cortical regions according to granularity indices across the cerebral cortex (as reported by von Economo (2009), further discussed by Beul and Hilgetag (2014), and made available by Scholtens et al. (2016)). Granularity indices: 0- allocortex (not a part of the neocortex), 1- agranular, 2- slightly granular, 3,4,5- increasingly granular, 6- granular. The entire atlas (2) and its components (3) can be seen from different viewpoints: A- top, B- occipital, C- lateral (left), D- bottom

We hypothesize that suitable clustering of the T1 layer composition across the entire cortex should correlate to spatially defined cortical regions with distinct cytoarchitectonic features. While many cortical atlases exist, the most robust reference atlas for cortical cytoarchitecture is the granularity atlas (von Economo, 2009). The granularity atlas divides each hemisphere into about a dozen continuous regions, each labelled with a granularity index 1-6 according to the level of overall cross section cellular granularity observed histologically (see figure 1, parts 2-3).

Analyzing patterns in this complex laminar dataset is no simple task, since the ability to visualize the entire dataset simultaneously is limited, and therefore accurate whole-brain clustering must be accomplished. The first challenge in clustering the dataset relates to its surface-based nature, in which the spatial locations of the data are not given across pixels or voxels, but rather across vertices on a triangulation surface. The second challenge relates to the dimensionality and low variability of the data, which includes six T1 layer widths with an overall average cortical cross section of approximately two millimeters. The third and final challenge relates to both spatial dispersion and multidimensionality: accurate clustering must take into consideration not only the T1 layer composition at each given datapoint, but also the compositions of its neighboring datapoints. The reasoning for these considerations is that cortical regions have distinct cytoarchitectonic features, based on the similarity of the laminar composition across them. Furthermore, at times even smaller regions vary in their cytoarchitectonic features. For example, gyral caps (peaks) and sulcal fundi (valleys) vary in their overall thickness as well as in their laminar substructure.

### 1.3 Existing methods for clustering multidimensional, graph-based datasets

One popular approach for clustering data with spatial information is by using graphs. In this approach, graph vertices represent data points in space and graph edges connect vertices that are close in space. Edges can be weighted reflecting the distance between the points. Over the years, a plethora of clustering algorithms have been developed for the problem, including some of the following: similarity graph connectivity clustering (Hartuv and Shamir, 2000), density-based clustering using both attribute similarity and spatial proximity (Liu et al., 2012), distributed K-means clustering of mesh networks (Ramesh, 2015), and community detection in networks (Girvan and Newman, 2002; Traag et al., 2019).

In medical imaging, various algorithmic approaches have been proposed for dealing with graph-based image segmentation (Chu et al., 2002) and density-based image segmentation using super-pixels (Zhang et al., 2017). In the field of MRI neuroimaging, different algorithmic approaches have been developed for threshold-free, surface-based clustering (Lett et al., 2017), as well as for community detection in functional networks (Akiki and Abdallah, 2019). Recently, a novel algorithm in the field of omics has been developed for clustering cell omics profiles using the spatial organization of the cells (Singhal et al., 2022). The algorithm, Building Aggregates with a Neighborhood Kernel and Spatial Yardstick (BANKSY), clusters multidimensional omics data across a surface-based spatial representation.

## 2. Methods and materials

### 2.1 Histological dataset- BigBrain segmentation

The histological dataset was provided by BigBrain, a high-resolution, three dimensional histological model of the human brain (Amunts et al., 2013). This dataset provides a three-dimensional segmentation of all cortical and laminar surfaces in the BigBrain, which have been segmented automatically based on histological intensities along cortical profiles sampled between the pial and white matters throughout the cortex. These surfaces were used to evaluate cortical thickness gradients and the contributions of different cortical lamina to these gradients (Wagstyl et al., 2020). Data is freely available at: https://bigbrainproject.org/

### 2.2 Neuroimaging datasets- MRI T1 layers

#### Human (N=33)

The human neuroimaging dataset includes (N=33) healthy human subjects, 16 male and 17 female, 18-78 years old, all neurologically and radiologically healthy with no history of neurological diseases. Most of the dataset includes thirty subjects (N=30) from the same dataset used by Shamir et al. (2022), and three additional subjects (N=3) were chosen as exemplary subjects from three groups of interest. All subjects signed informed consent before enrollment in the study. The imaging protocol was approved by the institutional review boards of Sheba Medical Center and Tel Aviv University, where the MRI investigations were performed. Each subject was then scanned in a 3T Magnetom Siemens Prisma (Siemens, Erlangen, Germany) scanner with a 64-channel RF coil and gradient strength of up to 80 mT/m at 200 m/T/s. The scans include the following sequences:

1. An MPRAGE sequence, with the following parameters: TR/TE = 1750/2.6 ms, TI = 900 ms, 1×1×1 mm^3^, 224×224×160 voxels, each voxel fitted with a single T1 value.
2. An inversion recovery echo planar imaging (IR EPI) sequence, with the following parameters: TR/TE = 10,000/30 ms and 60 inversion times spread between 50 ms up to 3,000 ms, 3×3×3 mm^3^, 68×68×42 voxels, each voxel fitted with up to 7 T1 values (Lifshits et al., 2018).

The first sequence was used as an anatomical reference, as well as for delineating the cortical surfaces, and the second sequence was used for characterizing the cortical layers. The acquired images were processed according to the framework for cortical laminar composition analysis (Shamir et al., 2019; Shamir and Assaf, 2021b; Shamir et al., 2022). The thirty-subject average dataset is free available at: https://github.com/ittais/Laminar_Connectivity

#### Macaque (N=1)

The macaque neuroimaging dataset is taken from Shamir and Assaf (2021b) and includes a single macaque brain that was obtained from the Mammalian MRI (MaMI) database (Assaf et al., 2020). No animals were deliberately euthanized for the present study. The excised macaque brain was formalin fixated, and some 24 h before MRI it was placed in phosphate-buffered saline for rehydration. For the scan, the brain was placed in a plastic bag and immersed in fluorinated oil (Flourinert, 3 M) to minimize image artifacts caused by magnet susceptibility effects. The brain was scanned on a 7 T/30 Bruker scanner with a 660 mT/m gradient system. The protocol was approved by the Tel Aviv University ethics committee on animal research. The scans include the following sequences:

1. A T1w sequence with a 3D modified driven equilibrium Fourier transform (MDEFT), with the following parameters: TR/TE = 1300/2.9 ms, TI = 400 ms, 0.2×0.2×0.2 mm^3^, 300×360×220 voxels.
2. An inversion recovery sequence using 3D FLASH, with the following parameters: TR/TE = 1300/4.672 ms and 44 inversion times spread between 25 ms up to 1,000 ms, each voxel fitted with up to 8 T1 values, voxel size 0.67×0.67×0.67 mm^3^, 96×96×68 voxels (similarly to Lifshits et al., 2018).

The first sequence was used as an anatomical reference with high gray/white matter contrast for segmentation and estimation of cortical surfaces (similar to the clinical MPRAGE sequence), and the second sequence was used for characterizing the cortical layers. The acquired images were processed according to the framework for cortical laminar composition analysis (Shamir et al., 2019; Shamir and Assaf, 2021b; Shamir et al., 2022).

### 2.3 Ground truth reference- granularity atlas

The cortical atlas of granularity indices (as shown in figure 1, parts 2-3) was used as the ground truth reference for cytoarchitecture. The atlas labels cortical regions according to the overall level of cytoarchitectonic granularity observed histologically across cortical cross sections (von Economo, 2009; Beul and Hilgetag, 2014; Scholtens et al., 2016). A similar atlas of cortical granularity was used for the macaque brain, based on a map of cytoarchitectonic features across the primate cortex (Beul and Hilgetag, 2019; Shamir and Assaf, 2021b).

### 2.4 BANKSY algorithm adaptation

In this study we use the BANKSY algorithm, originally developed in the field of omics for spatial clustering genomic and other biological data (Singhal et al., 2022). We adapt and implement the algorithm on surface-based cortical laminar composition datasets, both histological (BigBrain) and neuroimaging (MRI T1 layers). Our implementation of the algorithm includes the following steps:

1. Neighbor-augmented matrix construction: For each hemisphere, we construct a neighbor-augmented matrix as described by Singhal et al. (2022) using cortical layer width values instead of omics expression values. The matrix B is M×N, where N≅150,000 columns, corresponding to vertices on the cortical surface, and M=12 (see equation 1). Each column in the top half of the matrix B_top_ includes the six cortical layer widths for the vertex, and the bottom half B_bottom_ includes the six layer widths averaged over the neighborhood of the vertex. The contribution of each component is controlled by parameter *λ* (see equations 1-3). The neighborhood values are calculated by locating the thirty nearest vertices on the cortical surface and averaging them using the inverse of the distance to the original vertex, after normalizing the sum of the distances to one (see equation 4 and 5).

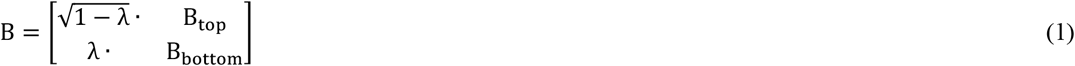 Where: *λ*-neighborhood weight, we used *λ* = 0.3 (Singhal et al. 2022)

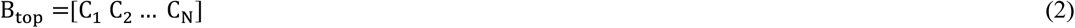 Where: C_i_- cortical layer widths for vertex i

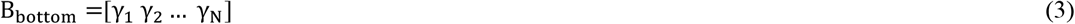 Where: λ_i_- average cortical layer widths of neighborhood of vertex i

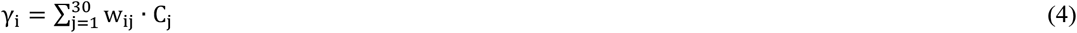 Where: C_j_- cortical layer widths for neighbor j of vertex i, w_ij_- weight for neighbor j

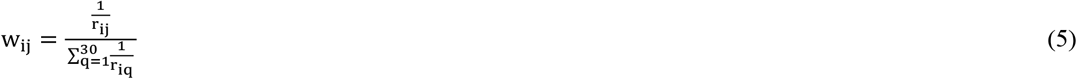 Where: r_ij_- distance of neighbor j from vertex i For the histological dataset, because of its assumed high resolution and accuracy, we only used the six layer width values for each vertex. In other words, the neighbor-augmented matrix was ~150,000× 12 as described above. For the neuroimaging datasets, because the T1 layers correspond to laminar components and lack a one-to-one correspondence to the histological cortical layers, we used the six layer width values as well as the overall cortical width values. The addition of the overall cortical width can be further explained by the relatively more precise measurement of the cortical cross section segmentation using MRI (Fischl, 2012). Overall, the neighbor-augmented matrix was ~150,000×14, including six cortical layer widths and the overall cortical width of the vertex, and six averaged cortical layer widths and the averaged overall cortical width of its neighborhood.
2. Neighbor-augmented matrix clustering: The neighbor-augmented matrices for both hemispheres are consolidated into a single whole-brain matrix for clustering (see equation 6).

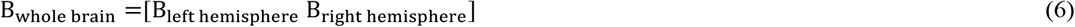 For both the histological and the neuroimaging datasets we clustered the corresponding matrices using a simple unsupervised K-means clustering algorithm, which partitions the vectors (~150,000×2 matrix columns) into K clusters. Aside from the variation in the construction of the neighbor-augmented vectors (see step 1), we used a different number of clusters K for the histological and neuroimaging datasets. For the histological dataset we got the best results when using K=6, and for the neuroimaging datasets (N=30 subjects) we repeatedly got the best results using K=4. For the first neuroimaging subject, the clusters were relabeled 1 to 4 in increasing levels of average granularity according to the granularity atlas (see figure 1, parts 2-3). To obtain consistent cluster labelling across subjects, the same K-means solution was used as seed for all other (N=29) subjects. For the macaque neuroimaging dataset we got the best results when using K=5.
3. Evaluating the clustering results: After assigning a label to each vertex, the clusters were plotted across the cortical surfaces and visually assessed for hemispheric symmetry and overall similarity to the granularity atlas. To integrate the results across subjects, we used the Brainnetome atlas, which partitions the cortex into 210 cortical regions (Fan et al., 2016). Three visual and quantitative assessments were then performed:
  a. Hemispheric symmetry:
    i. A cross-subject cluster label was assigned to each cortical region according to a majority vote of all vertices in that region across all subjects (N=30). To visually assess the symmetry, the clusters were plotted across the cortical surfaces.
    ii. To quantitatively assess the level of symmetry, for each subject, each region was assigned a per-subject label according to majority vote. The percentage of regions that differ in their majority vote clusters between the hemispheres was calculated for all (N=30) subjects and compared to randomly spatially permuted datasets.
  b. Inter-subject variability: To assess inter-subject variability, the per-subject region labels described in a(ii) were used. The standard deviation in cluster assignment per region across all (N=30) subjects was then measured and plotted across the cortical surfaces.
  c. Similarity to granularity atlas: To visually assess the similarity, the cross-subject region labels described in a(i) were used. The clusters were plotted across the cortical surfaces and compared to a simplified version of the granularity atlas. To quantitatively assess the similarity, a hypergeometric test was performed comparing the resulting clusters to the simplified groups of granularity indices. The number of regions in each index and in each cluster was computed, and the significance of the overlap between each index-pair was computed. P-values were corrected using the Bonferroni correction for multiple comparisons.

## 3. Results

### 3.1 Clustering the histological dataset

K-means clustering of the neighbor-augmented matrix for the histological dataset resulted in six distinct clusters that show correspondence to most of the six granularity indices in the granularity atlas (see results in figure 2). When examining the resulting six clusters, it appears that the methodology successfully delineated occipital regions with increasingly high granularity, alongside successful delineation of temporal regions with mid-to-low granularity levels (figure 2, row 1B and C, respectively). Overall visual assessment of the clusters shows high hemispheric symmetry between the left and right hemispheres. When examining the six clusters individually, additional interesting features appear. As seen, the process successfully delineated the allocortex (figure 2, row 2). This result is expected given the fact that no layer data is included for the allocortex, which is not a part of the neocortex and has no granularity index labelling. The other clusters seem to correspond to regions with increasing granularity indices, from more frontal regions that are considered more agranular (figure 2, rows 3 and 4), to less frontal regions that are considered more granular (figure 2, rows 5 and 6), up to occipital regions that are considered entirely granular (figure 2, row 7). Furthermore, the clustering process seems to have accurately delineated the postcentral gyrus as part of the less granular cortex (figure 2, row 5A) and the precentral gyrus as part of the granular cortex (figure 2, row 7A). A notable feature across all individual clusters is the differentiation across the cortical folding, i.e., gyral caps and sulcal fundi, which are known to differ in both thickness and laminar composition (MacDonald et al., 2000; Wagstyl et al., 2020).

**Figure 2:**
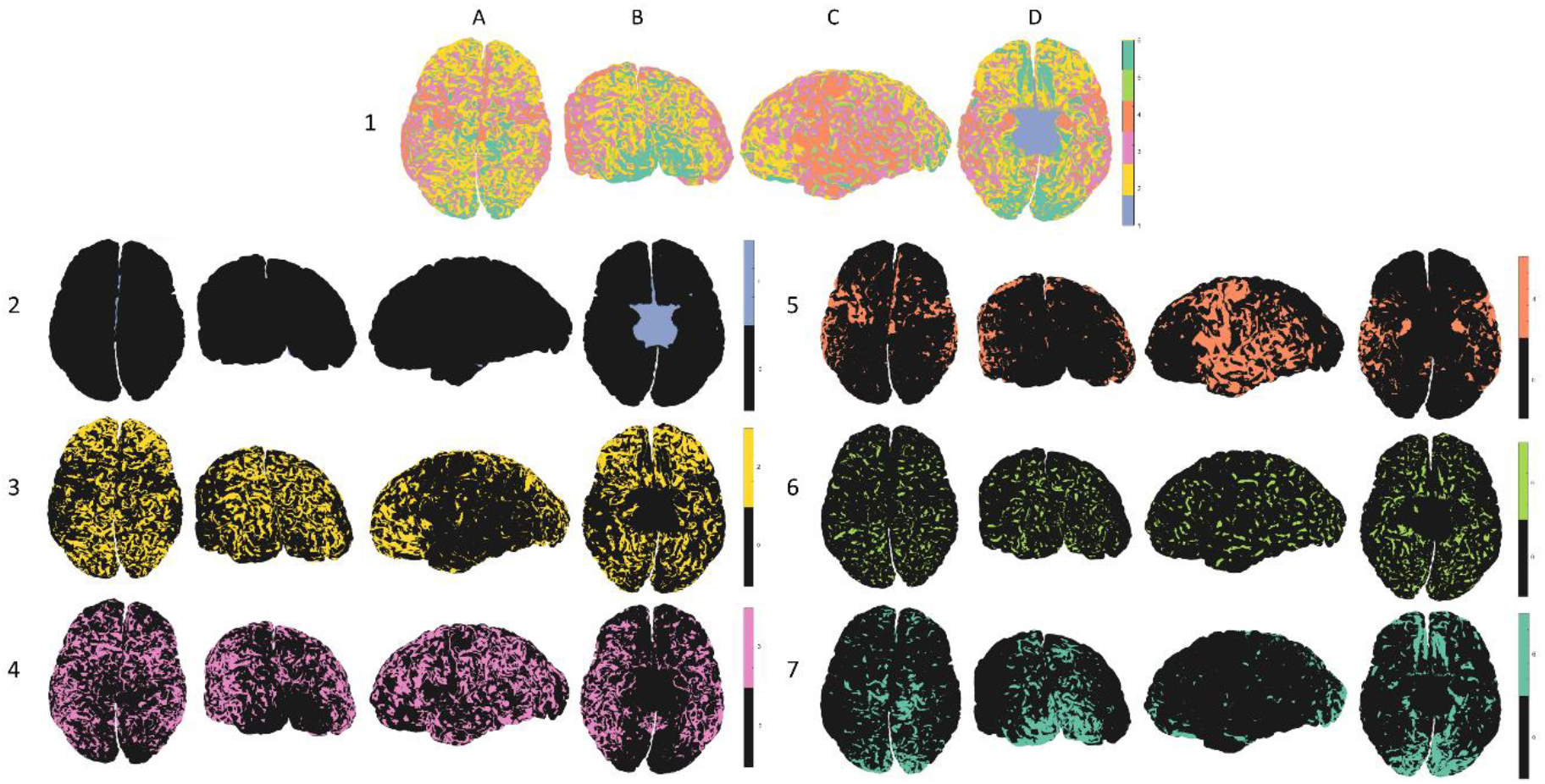
Clustering histological dataset (BigBrain): All six clusters can be seen in the top row (1), and each individual cluster can be seen on the rows below (2-7): 2- allocortex, 3,4,5-agranular to increasingly granular cortices, 6,7- increasingly granular to granular cortex. The clusters can be seen from different viewpoints: A- top, B- occipital, C- lateral (left), D- bottom

The effective clustering of cortical layers in the histological dataset, accurately identifying cortical regions with varying cytoarchitectonic features, provided a proof of concept for the potential applicability of the BANKSY clustering algorithm for our T1 layer neuroimaging datasets.

### 3.2 Clustering the neuroimaging datasets

The results of K-means clustering of the neighbor-augmented matrix for the neuroimaging dataset with K=4 clusters show correspondence to a coarser division of the granularity indices in the granularity atlas (see figure 3). When examining all four clusters simultaneously, once again it appears that the process successfully delineated occipital regions with increasingly high granularity, alongside successful delineation of temporal regions with mixed granularity levels (figure 3, row 1B and C, respectively). Additionally, relatively high hemispheric symmetry can be seen from a visual assessment of the clusters. When examining the four clusters individually, some interesting features appear. As seen, the process delineated the allocortex but merged it with the agranular cortex (figure 3, row 2). Unlike the histological dataset, the neuroimaging datasets do include layer information for the allocortex, which is not part of the neocortex but is in fact characterized by a laminar composition of three to four layers. The other clusters seem to correspond to a coarser division into regions with increasing granularity indices, from more frontal regions that are considered less granular (figure 3, row 3), to less frontal regions that are considered more granular (figure 3, row 4), ending once more in occipital regions with high granularity (figure 3, row 5). Accurate delineation of the precentral gyrus as part of the granular cortex is also apparent (figure 3, row 5A).

**Figure 3:**
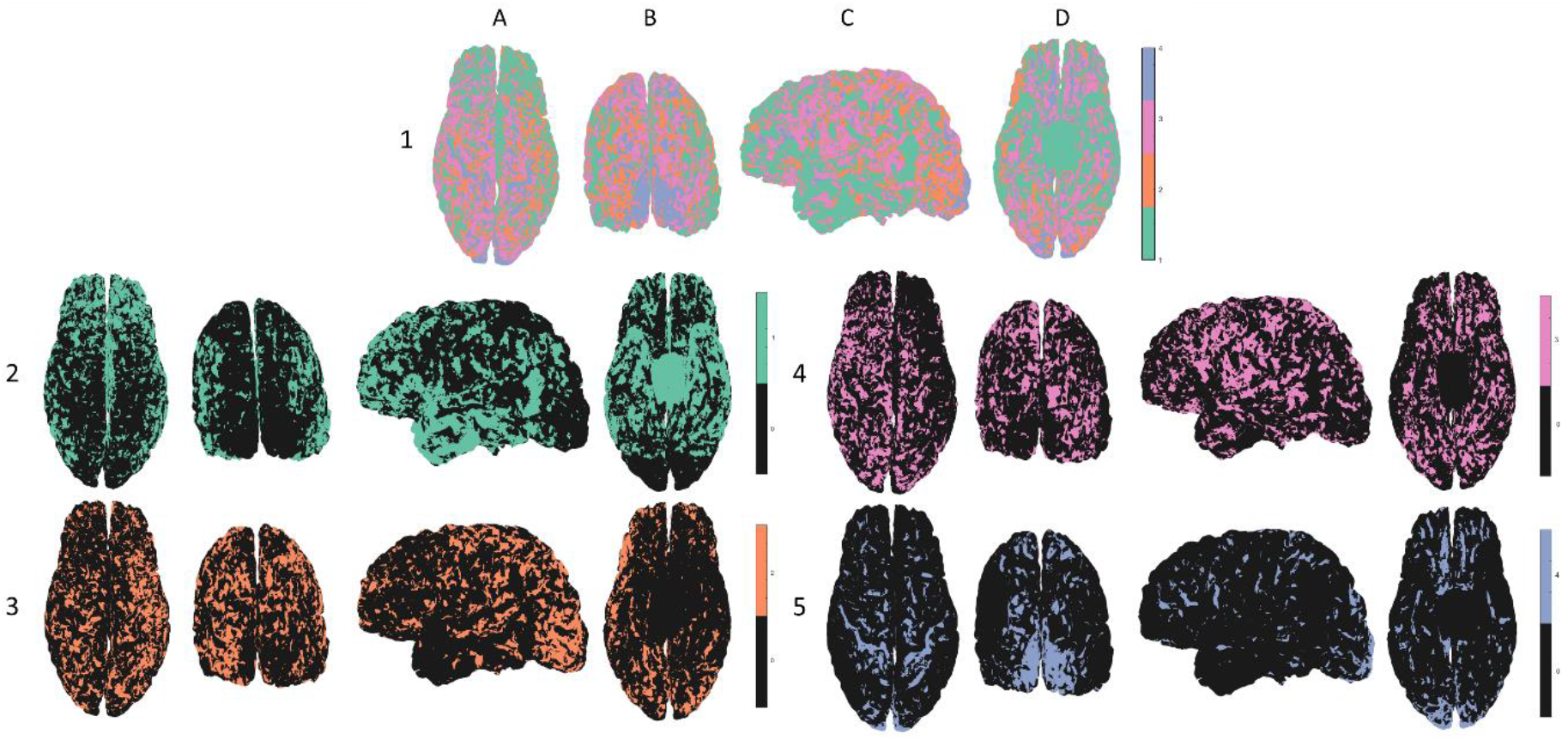
Clustering neuroimaging dataset for a single subject (MRI T1 layers): All four clusters can be seen in the top row (1), and each individual cluster can be seen on the rows below (2-5): 2- allocortex, agranular cortex, 3,4- increasingly granular, 5- granular cortex. The clusters can be seen from different viewpoints: A- top, B- occipital, C- lateral (left), D- bottom

Similar patterns were observed in the clustering results of all (N=30) subjects. The differentiation between cytoarchitecture in gyri and sulci that was previously observed for the histological dataset is also noticeable here across all individual clusters (see figure 4, part 1). Furthermore, when we randomly permute the spatial locations of the neighbor-augmented vectors in the neuroimaging dataset and the apply the same adaptation of the BANKSY algorithm, any delineation of regions of cytoarchitectonic importance disappears (see figure 4, part 2).

**Figure 4:**
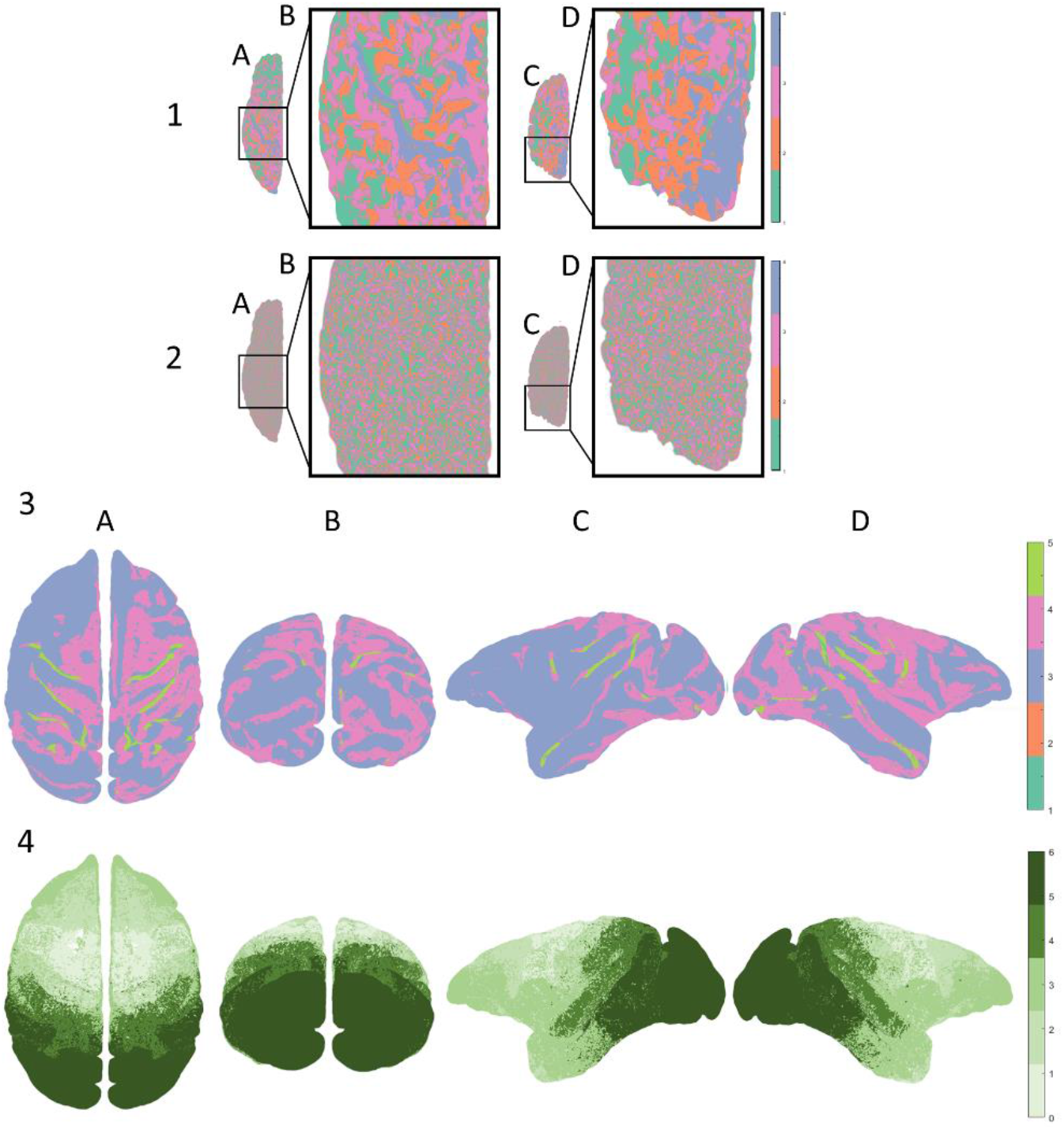
Clustering different neuroimaging datasets: Differences in clusters between an original and a randomized neuroimaging dataset (human): 1- Original dataset: the resulting clusters in a neuroimaging dataset showcase a differentiation between gyri and sulci across the cortical folding 2- Randomized dataset: the same neuroimaging dataset was permuted spatially and then clustered using the same algorithm, resulting in no delineation of any regions of cytoarchitectonic importance The results across the left hemisphere can be seen from a top view (A) of the precentral gyrus (B), and from an occipital view (C) of the primary visual cortex (D) 3- Clustering macaque neuroimaging dataset: all five resulting clusters can be seen from different viewpoints: A- top, B- occipital, C- lateral (left), D- lateral (right) 4- Granularity atlas (reduced): the cytoarchitectonic atlas of granularity indices (as seen in figure 1), adapted for the macaque brain (Beul and Hilgetag, 2019; Shamir and Assaf, 2021b) and reduced to five components

When examining the six resulting clusters for the excised Macaque brain, a similar cytoarchitectonic differentiation between gyri and sulci appears (see figure 4, part 3). While the excised macaque brain displays fewer dominant clusters, presumably relating to the formalin fixation, the resulting clusters display high hemispheric symmetry and a clear delineation of regions relating to the macaque motor cortex.

To further evaluate the performance of the algorithm on the neuroimaging datasets, we computed per-subject cluster assignment to each region using majority voting and measured the standard deviation of results across all (N=30) subjects. The regional majority vote was also used for measuring the overall accuracy of the results on a regional level, by finding cross-subject cluster assignment to each region using majority vote of the points in the region across all subjects (see figure 5, part 1). Standard deviation was used for measuring variability in per-subject regional cluster assignment across subjects (see figure 5, part 2). The evaluation reestablishes the coarser delineation of overall granularity patterns: high granularity in occipital regions and in the precentral gyrus, mixed granularity in temporal regions, low granularity in frontal regions (merged with the allocortex) and increasing granularity in less frontal regions. To quantitatively assess the overall similarity to the granularity atlas, we measured hypergeometric scores, testing the correspondence between the four clusters and four groups of granularity indices. Significant correspondence was found between cluster 1 and granularity indices 2-3, cluster 2 and indices 0-1, cluster 3 and index 4, and between cluster 4 and indices 5-6 (see figure 5, part 4).

**Figure 5:**
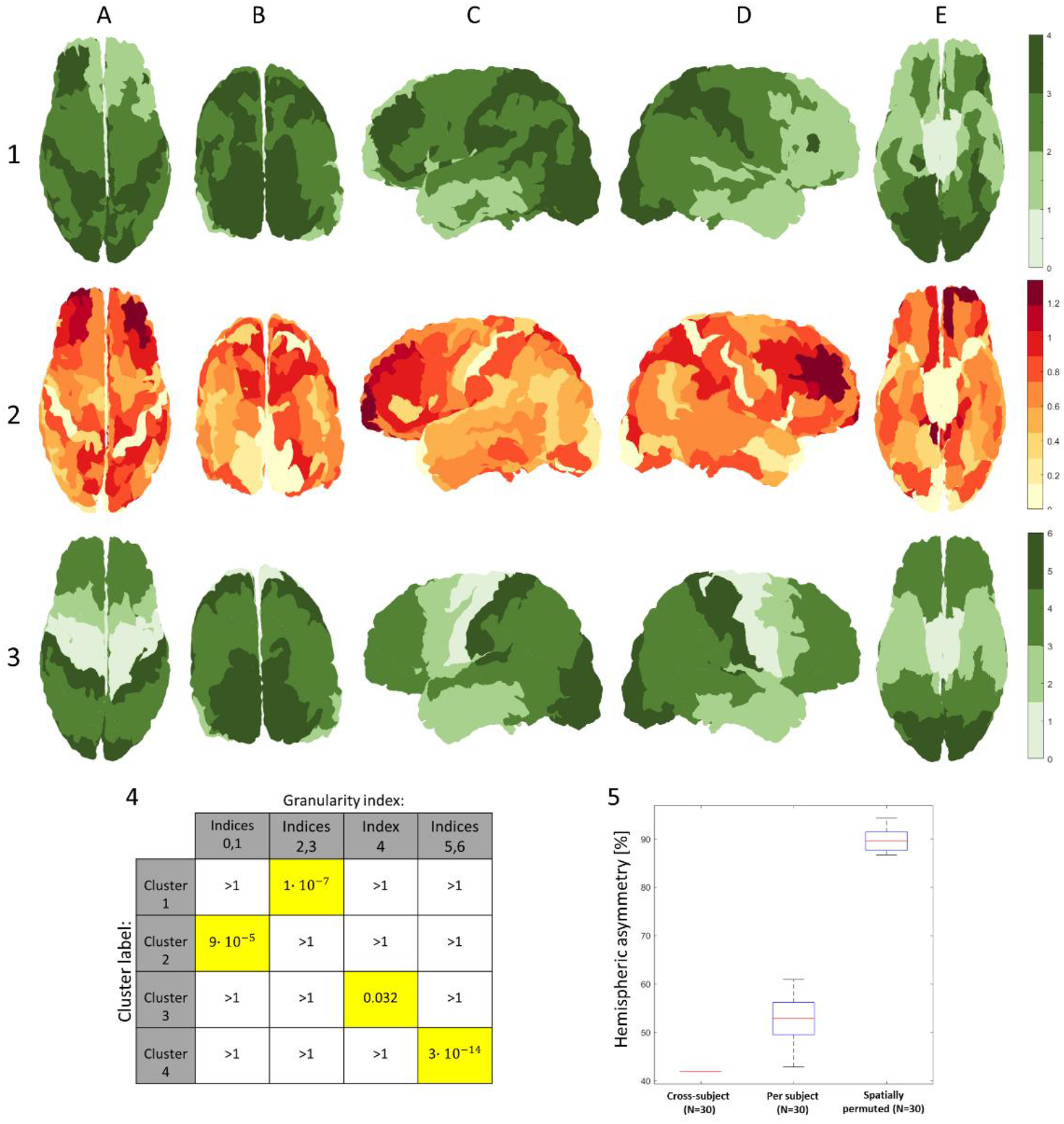
Inter-subject (N=30) clustering results and quantitative assessments: 1- Inter-subject majority vote: the most common cluster assigned to vertices belonging to each corresponding Brainnetome atlas region, evaluated across all subjects (N=30) 2- Inter-subject variability: the standard deviation of the majority vote clusters across Brainnetome atlas regions, evaluated across all subjects (N=30) 3- Granularity atlas (reduced): the cytoarchitectonic atlas of granularity indices (as seen in figure 1), reduced from the original six components to a coarser division including four components. Images are shown from different viewpoints: A- top, B- occipital, C- lateral (left), D- lateral (right), E-bottom 4- Similarity to granularity atlas: hypergeometric p-values for the correspondence between the four cross-subject clusters and four groups of granularity indices. P-values were Bonferroni-corrected for multiple comparisons. Significantly correlated pairs (p-value<0.05) are marked in yellow 5- Hemispheric asymmetry: boxplots of the distribution of the fraction of pairs of corresponding cortical regions (left and right hemispheres, N=105 pairs) that differ in their cluster assignment. Three distributions are shown: cross-subject region labels, when taking majority per region separately for each subject (N=30), and when repeating the latter for randomly spatially permuted datasets (N=30)

When assessing hemispheric symmetry, some asymmetry appears, particularly in regions with mixed granularity indices, such as frontal and temporal regions, which also exhibit high inter-subject variability. To quantitatively assess the hemispheric asymmetry, we measured the percentage of pairs of corresponding regions (mirroring regions between hemispheres, N=105) that differ in their cluster assignment. The datasets include the following three categories: cross-subject region labels, per-subject region labels (N=30), and per-subject region labels on randomly spatially permuted datasets (N=30). The results show high symmetry values for the original datasets in comparison to the spatially permuted datasets (see figure 5, part 5).

To examine subject-specific cytoarchitectonic features, we chose three additional exemplary subjects (N=3) from three groups of interest: a professional athlete, a professional musician, and a multilingual subject (polyglot). We applied the same clustering methodology on each subject and presented the clustering results in relation to the thirty-subject majority vote (see figure 6). The results show several notable features: the professional athlete exhibits relatively higher granularity in motor and premotor regions, as well as frontal regions (see figure 6, part 1), the professional musician exhibits higher granularity in motor and auditory regions (see figure 6, part 2), and the polyglot exhibits higher granularity in regions associated with language perception and formation (see figure 6, part 3). Subject handedness is also apparent in the results: both the athlete and the musician are right handed, and accordingly they exhibit higher granularity in motor regions of the left hemisphere, while the polyglot is left handed and accordingly exhibits high granularity in mirrored language regions of the right hemisphere.

**Figure 6:**
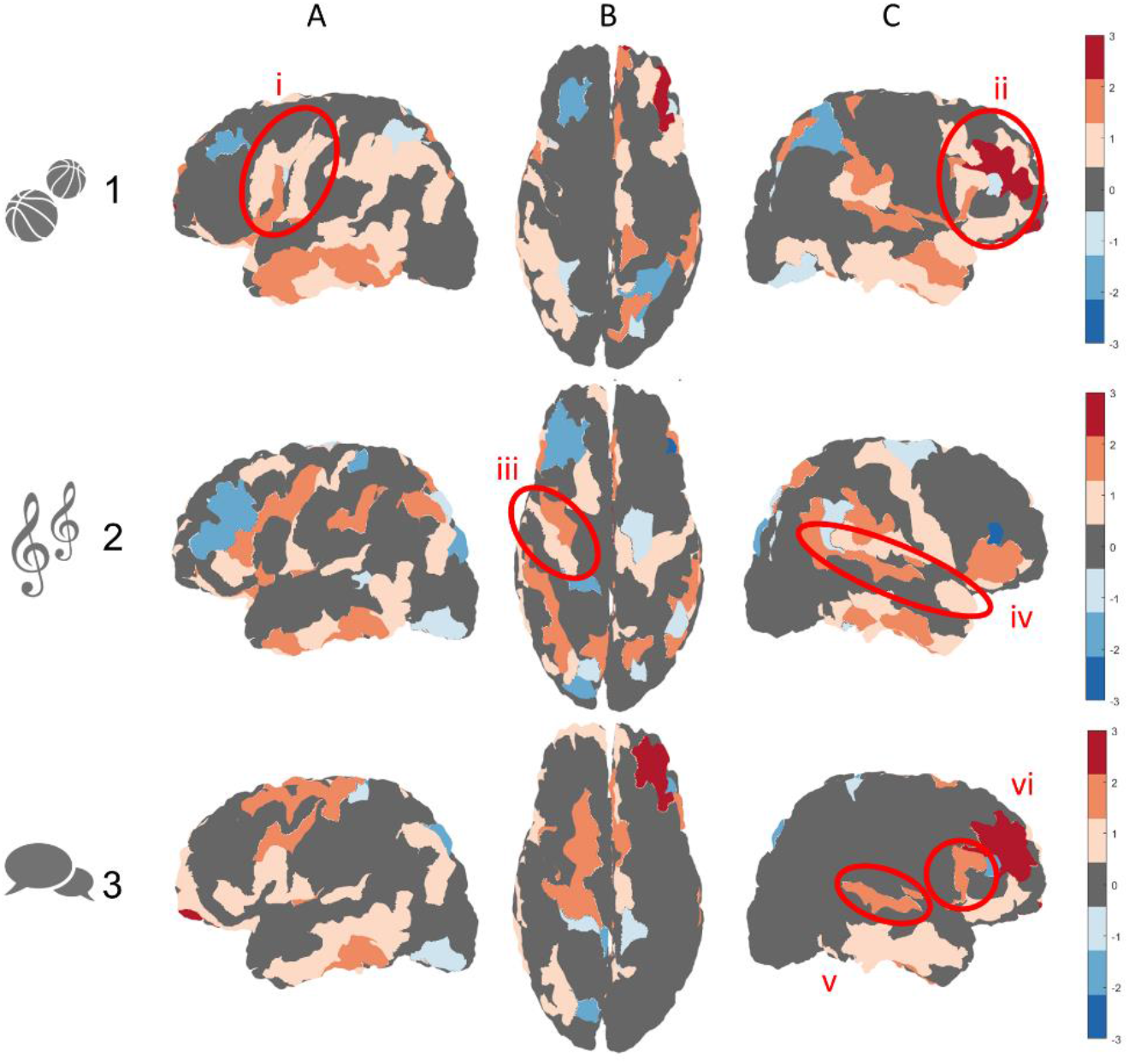
Relative clustering results for exemplary subjects (N=3) from different groups of interest: Clustering results are shown in relation to the thirty subject (N=30) majority vote (presented in figure 5, part 1), where regions with relatively higher granularity are presented in hot colors and regions with lower granularity are presented in cold colors: 1- A professional athlete (N=1): relatively higher granularity in motor and premotor regions (i), as well as frontal regions (ii) 2- A professional musician (N=1): relatively higher granularity in motor (iii) and auditory (iv) regions 3- A polyglot, or person with fluency in multiple languages (N=1): relatively higher granularity in regions associated with language perception and formation (v and vi) It should be noted that while the professional athlete and musician are both right handed, the polyglot is left handed. Images are shown from different viewpoints: A- lateral (left), B- top, C- lateral (right)

## 4. Discussion

In this study we cluster microstructural multilayered surface-based data in the cerebral cortex using adaptations of an omics algorithm called Building Aggregates with a Neighborhood Kernel and Spatial Yardstick (BANKSY). This algorithm was developed in the field of omics for clustering genomic and biological datasets based on both cell types and tissue domains and it has been shown to outperform related clustering methods for multiple types of datasets, including successful delineation of the cortical layers by clustering omics data from the dorsolateral prefrontal cortex (Singhal et al., 2022).

We first adapted the algorithm for the BigBrain histological dataset (Amunts et al., 2013), a high-resolution three-dimensional segmentation of all cortical and laminar surfaces in the brain (Wagstyl et al., 2020). This initial adaptation involved the use of the six width values of the laminar components and clustering into K=6 clusters, resulting in delineation of multiple cortical regions with distinct cytoarchitectonic features, including the allocortex, regions with increasing granularity from frontal to less frontal regions, and high granularity regions in the occipital cortex as well as the precentral gyrus. This adaptation provided an initial proof of concept for the applicability of the BANKSY methodology for other surface-based cytoarchitectonic datasets.

We then adapted the algorithm for the neuroimaging datasets, which include the laminar composition of six T1 layers across the cortical surfaces of (N=30) healthy subjects (Shamir et al., 2022). For this adaptation of the algorithm, we achieved optimal results when using the widths of the six laminar components combined with the overall cortical width, all clustered into K=4 clusters. The use of fewer clusters when clustering the neuroimaging datasets can be explained by the relatively lower resolution of these datasets compared to the ultrahigh resolution of the BigBrain histological dataset. The relatively lower resolution of the neuroimaging dataset can also account for the inclusion of the overall cortical width as input, alongside the more well-established and well-documented nature of MRI segmentation of overall cortical thickness. Once again, the clustering resulted in delineation of multiple regions with distinct cytoarchitectonic features, including regions with increasing granularity from frontal to less frontal regions, and high granularity in occipital regions as well as in the precentral gyrus.

The resulting clusters in both adaptations of the algorithm are characterized by a “patchy” appearance, compared to the more uniform nature of our chosen reference atlas of cytoarchitectonic features (von Economo, 2009). The “patchiness” appears to relate to a differentiation between gyri and sulci across the cortical folding, known to vary in both overall thickness as well as in laminar composition (MacDonald et al., 2000; Wagstyl et al., 2020). This explanation is strengthened by the fact that this feature appears in both the histological dataset, which includes only laminar composition as input, as well as in the neuroimaging datasets, which also incorporates the overall cortical thickness as input. The difference in uniformity across cortical regions can also be attributed to methodological differences relating to the labeling process of the granularity atlas. This atlas is the result of a manual labeling process by histologists almost a century ago according to cellular features observed across entire cortical regions. By comparison, our clustering methodology involves unsupervised labeling of present-day datasets according to assumed cellular features on a vertex-wise basis. The vertex-wise clustering better suits the nature of the neuroimaging datasets and the regional variability in laminar composition. Nevertheless, to better assess the accuracy of the results quantitatively we used a majority vote for clusters across cortical regions. Our analysis found a significant correspondence between each of the four resulting clusters and a different set of granularity indices. Furthermore, examination of the clustering results for three exemplary subjects with unique skills highlights the applicability of this framework in the exploration of the structural mechanisms and cytoarchitectonic features behind different skillsets.

The results of this adaptation of the BANKSY algorithm for clustering of multilayered laminar datasets across cortical surfaces in the brain further highlights the role of MRI neuroimaging as a probe of tissue cytoarchitecture. The correspondence between T1 layer clusters and regions with distinct cytoarchitectonic features provides a robust validation of the T1 imaging framework for cortical laminar composition analysis (Shamir et al., 2019; Shamir and Assaf, 2021a, 2021b; Shamir et al., 2022). In the future, the clustering methodology presented here can be implemented on large-scale groups of subjects not only to create a new and updated in-vivo atlas of cytoarchitectonic features, but also to further characterize subject-specific features associated with various abilities and skills.

